# Physics-Informed Neural Networks for Parameter Recovery in the Repressilator Oscillatory Model

**DOI:** 10.64898/2026.05.12.724679

**Authors:** Bernat Casajuana, Roger Casals-Franch, Adrián López García de Lomana, Pere Martí-Puig, Jordi Villà-Freixa

**Affiliations:** Computational Biochemistry and Biophysics Lab, Bioinformatics and Bioimaging (BI-SQUARED) Research Group, Department of Biosciences, Universitat de Vic - Universitat Central de Catalunya (UVic-UCC), 08500 Vic, Spain; Institut de Recerca i Innovació en Ciències de la Vida i de la Salut a la Catalunya Central (IRIS-CC), 08500 Vic, Spain; BioMedical Center, School of Health Sciences, University of Iceland, 102 Reykjavik, Iceland; Data and Signal Processing (TDS) Research Group, Faculty of Sciences, Technology and Engineering, Universitat de Vic - Universitat Central de Catalunya (UVic-UCC), 08500 Vic, Spain

**Keywords:** Physics-informed neural networks, Repressilator, Parameter estimation, Systems biology, Inverse problems

## Abstract

Parameter estimation in nonlinear biological dynamical systems is a difficult inverse problem because the governing equations are often stiff or oscillatory, the data are sparse and noisy, and the objective landscape is non-convex. Physics-informed neural networks (PINNs) offer an alternative to purely simulation-based calibration by representing state trajectories with neural networks while penalizing violations of the governing equations. This paper studies the empirical reliability of PINNs for recovering the parameters of the repressilator, a synthetic genetic oscillator formed by three cyclically repressive genes. We use synthetic time-series generated from the standard ordinary differential equation model and train inverse PINNs to estimate the production parameter *β* and the Hill coefficient *n*. The study varies observation noise, partial observation of repressors, sampling density, sensitivity to initial parameter guesses, and the difference between stable and oscillatory regimes. The results show that PINNs can reconstruct trajectories accurately when the model structure is correct and the three repressors are observed, but parameter recovery is more fragile than trajectory fitting. Noise, sparse sampling, unobserved variables, and unfavorable initial guesses increase the risk of biased estimates. The stable regime is easier to reconstruct, whereas the oscillatory regime provides richer information but also exposes optimization sensitivity. These findings support PINNs as a useful reverse-engineering tool for small gene-regulatory ODE models, while highlighting the need for repeated runs, uncertainty reporting, and experimental designs that improve identifiability.

## 1 Introduction

Mathematical models of gene regulation help connect molecular interactions with dynamical phenotypes such as adaptation, bistability, and oscillation. In systems biology, these models are commonly expressed as ordinary differential equations (ODEs) whose parameters summarize production rates, degradation rates, binding cooperativity, and regulatory thresholds. Stochastic formulations are also important when copy-number fluctuations or cell-to-cell variability shape the observed phenotype, as shown for the genetic toggle switch and for evolved noisy oscillatory genetic networks [16, 6]. Estimating parameters from data is a central reverse-engineering task, but it remains challenging because biological time series are often short, noisy, partially observed, and generated by nonlinear systems with correlated or weakly identifiable parameters [5, 7, 4, 9].

Classical approaches rely on repeated numerical simulation and optimization. Global and hybrid metaheuristics have improved robustness in biological calibration problems [12, 10], while regularized and scalable nonlinear programming formulations address overfitting, local minima, and larger model classes [14, 9]. A complementary line of work addresses structural rather than parametric uncertainty. When an ODE model cannot explain the observed data, topological augmentation reformulates it as a stochastic differential equation and uses the residual stochastic terms to locate missing interactions and guide iterative model improvement [15]. Nevertheless, even small oscillatory systems can be difficult because matching an observed trajectory does not guarantee that the recovered parameters are unique or biologically meaningful.

Physics-informed neural networks (PINNs) provide a complementary route for solving forward and inverse problems governed by differential equations [11]. A neural network represents the unknown state trajectory, automatic differentiation evaluates time derivatives, and the loss function combines data mismatch, initial or boundary conditions, and residuals of the governing equations. In inverse settings, model parameters are optimized together with the neural-network weights. PINNs have become attractive in scientific machine learning because they can combine scattered data with mechanistic constraints and can be implemented without repeatedly calling an external ODE solver [8].

This work evaluates PINNs on the repressilator, a canonical synthetic gene oscillator introduced by Elowitz and Leibler [3]. The repressilator is small enough to allow controlled experiments, but it is dynamically rich enough to expose practical difficulties in parameter recovery, including oscillations, phase shifts, hidden variables, and sensitivity to the Hill coefficient.

We use simulated protein-concentration time series to characterize when an inverse PINN can recover the repressilator parameters. The experiments vary observation noise, the number of measured repressors, temporal sampling density, initial guesses for the unknown parameters, and the dynamical regime. These factors were chosen because they separate state reconstruction from parameter identification. The ODE residual is expected to regularize trajectory learning under moderate noise or sparse data, while accurate recovery of *β* and *n* still requires observations that carry enough information about amplitude, phase, and nonlinear repression. Recovery should therefore deteriorate when repressors are unobserved, when sampling misses informative oscillatory phases, or when optimization begins far from a physically meaningful basin.

## 2 Methods

### 2.1 Repressilator Model

We use the standard dimensionless three-protein repressilator model:

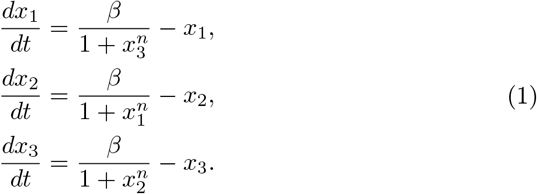

Here *x*_1_, *x*_2_, and *x*_3_ are repressor concentrations, *β* controls maximal production, and *n* is the Hill coefficient. The degradation coefficient is fixed by nondimensionalization. The model was simulated with initial condition (1.0, 1.0, 1.2) over *t* ∈ [0, 20] using 1000 time points. The main oscillatory setting used *β* = 5.0 and *n* = 3.0; a stable comparison regime used *β* = 5.0 and *n* = 1.5.

### 2.2 Synthetic Data

Synthetic observations were generated by numerical integration of Eq. 1. Gaussian noise was added to each state:

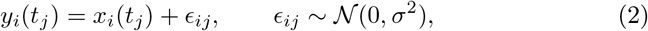

where *σ* is set as a fraction of the mean peak-to-peak signal amplitude. The codebase cloned from the project repository [2] contains 100 pre-generated datasets. These sweep four values of *β*, five values of *n*, and five noise levels. The experiments reported in this article use in-memory datasets generated by the same code path to allow repeated seeds and matched configurations.

### 2.3 Inverse PINN Formulation

The inverse PINN represents (*x*_1_, *x*_2_, *x*_3_) with a fully connected neural network *u*_*θ*_(*t*). The vector *θ* denotes the trainable coefficients and biases in the network layers. These network weights define the shape of the approximated trajectory and are distinct from the scalar loss weights *λ*_*f*_, *λ*_0_, and *λ*_*y*_ used to balance the residual, initial-condition, and observation terms. The physical parameters 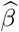 and 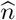 are optimized at the same time as *θ*. Automatic differentiation gives *du*_*θ*_*/dt*, and the residuals are

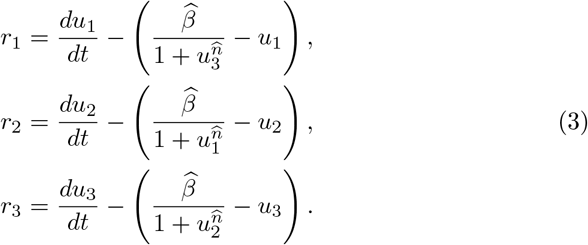

The training objective is

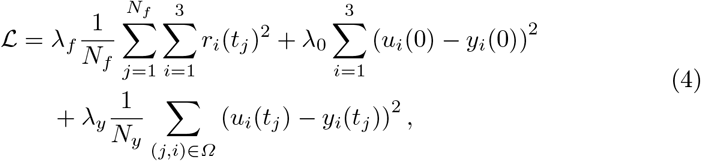

where *Ω* indexes the observed state components and observation times. In the implementation, the network has five hidden layers of 100 neurons, sinusoidal activations, Glorot initialization, and a softplus output transform to enforce positive concentrations. Models are trained with Adam using DeepXDE with TensorFlow backend [8]. The current experiment drivers use 3000 iterations for the sampling, partial-observation, and initial-guess studies, and 5000 iterations for the noise and regime-comparison studies.

### 2.4 Experimental Design

Five experiments were implemented as independent drivers in the companion repository. The current executable scripts use compact sweeps suitable for repeated local or cluster runs, while preserving the main stress tests needed for this paper. Each run records the relative error in *β*, the relative error in *n*, their arithmetic mean, and the state-reconstruction RMSE against the clean simulation. For parameter *p*∈ {*β, n*}, the relative error is 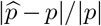 . The trajectory error is computed over the full clean trajectory, not only over the observed points,

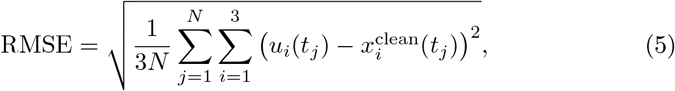

where *N* = 1000 for the simulated time grid. When an experiment has repeated seeds, the reported summaries use the mean and standard deviation across runs.

Table 1 summarizes the role of each driver. The first three experiments perturb the data available to the PINN by changing noise, observability, and sampling. The fourth experiment keeps the data fixed and changes only the optimizer initialization for the physical parameters. The final experiment compares a stable trajectory with an oscillatory one, so that reconstruction difficulty and parameter informativeness can be assessed under different dynamics.

**Table 1.**
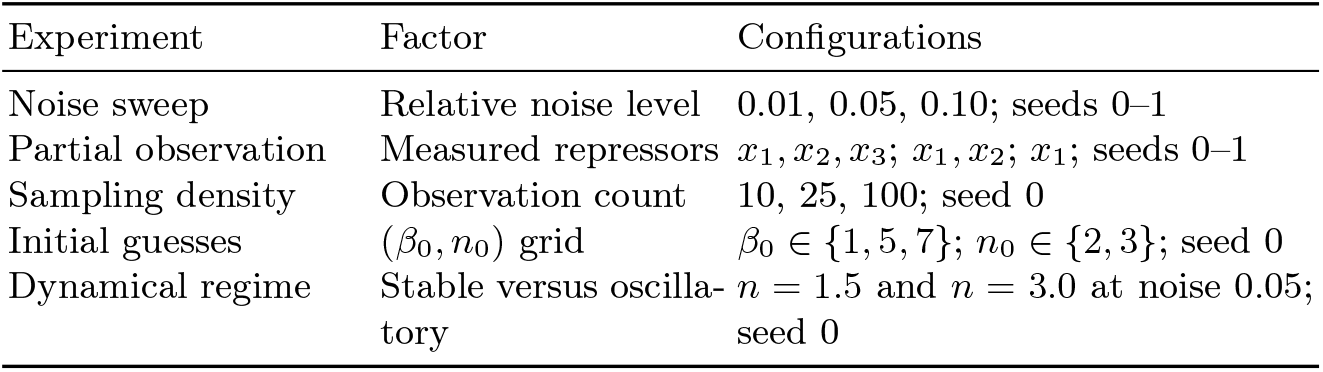
Empirical factors evaluated by the inverse-PINN experiment drivers. All runs use *β* = 5.0 and *n* = 3.0 unless the regime comparison explicitly sets *n* = 1.5 for the stable case.

| Experiment | Factor | Configurations |
| --- | --- | --- |
| Noise sweep | Relative noise level | 0.01, 0.05, 0.10; seeds 0–1 |
| Partial observation | Measured repressors | $x_1, x_2, x_3$; $x_1, x_2$; $x_1$; seeds 0–1 |
| Sampling density | Observation count | 10, 25, 100; seed 0 |
| Initial guesses | $(\beta_0, n_0)$ grid | $\beta_0 \in \{1, 5, 7\}$; $n_0 \in \{2, 3\}$; seed 0 |
| Dynamical regime | Stable versus oscillatory | $n = 1.5$ and $n = 3.0$ at noise 0.05; seed 0 |

## 3 Results

### 3.1 Forward Reconstruction in Stable and Oscillatory Regimes

The first diagnostic compares ODE-generated trajectories with PINN predictions in stable and oscillatory regimes. In the stable setting, the network accurately follows the relaxation to a common steady state for all three repressors (Fig. 1). This regime is comparatively easy because the long-time data contain a dominant equilibrium signal and the phase information is not critical.

**Fig. 1.**
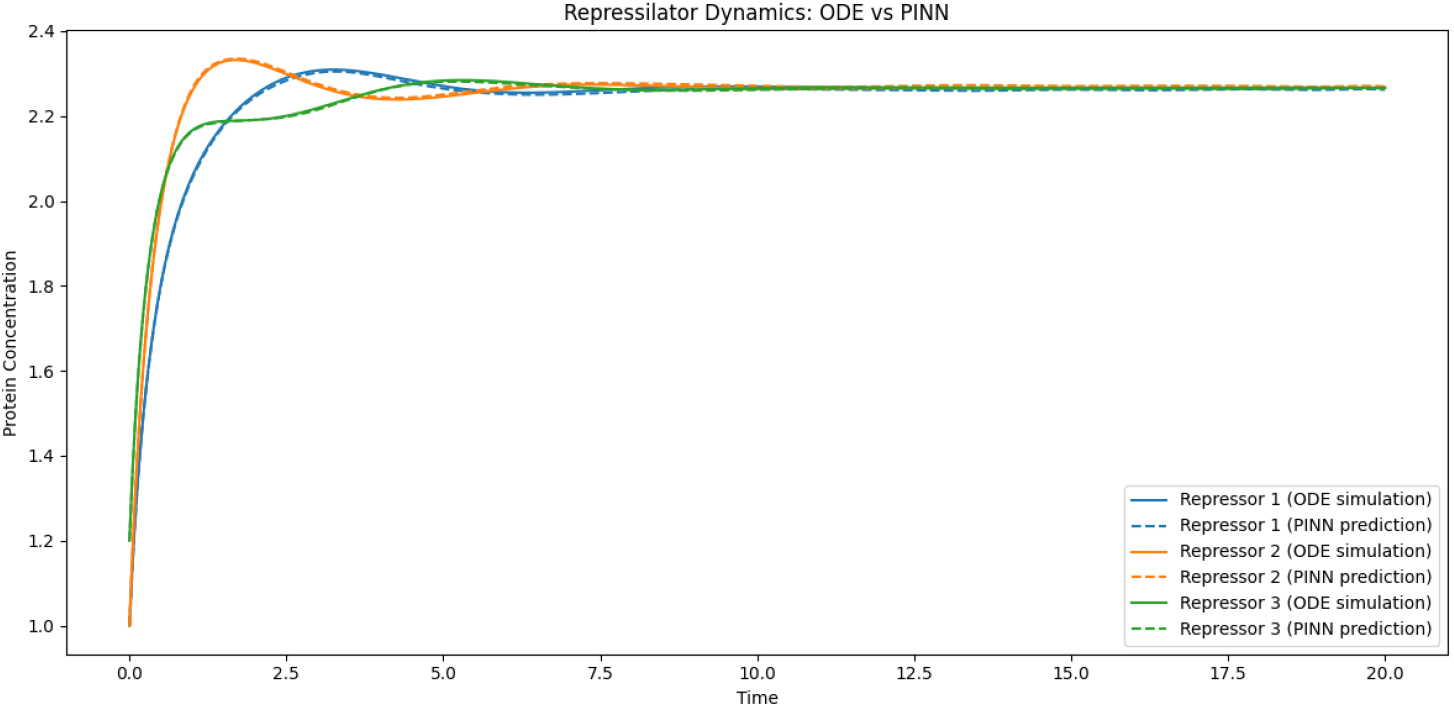
PINN reconstruction of a stable repressilator trajectory. Solid lines are ODE simulations and dashed lines are PINN predictions.

In the oscillatory setting, the PINN also captures the qualitative phase-shifted dynamics of the three repressors (Fig. 2). The match is strongest where the observations constrain peaks and troughs. Small deviations around rapidly changing phases are more consequential for parameter inference than for visual trajectory reconstruction, because *β* and *n* control amplitude, nonlinearity, and timing simultaneously.

**Fig. 2.**
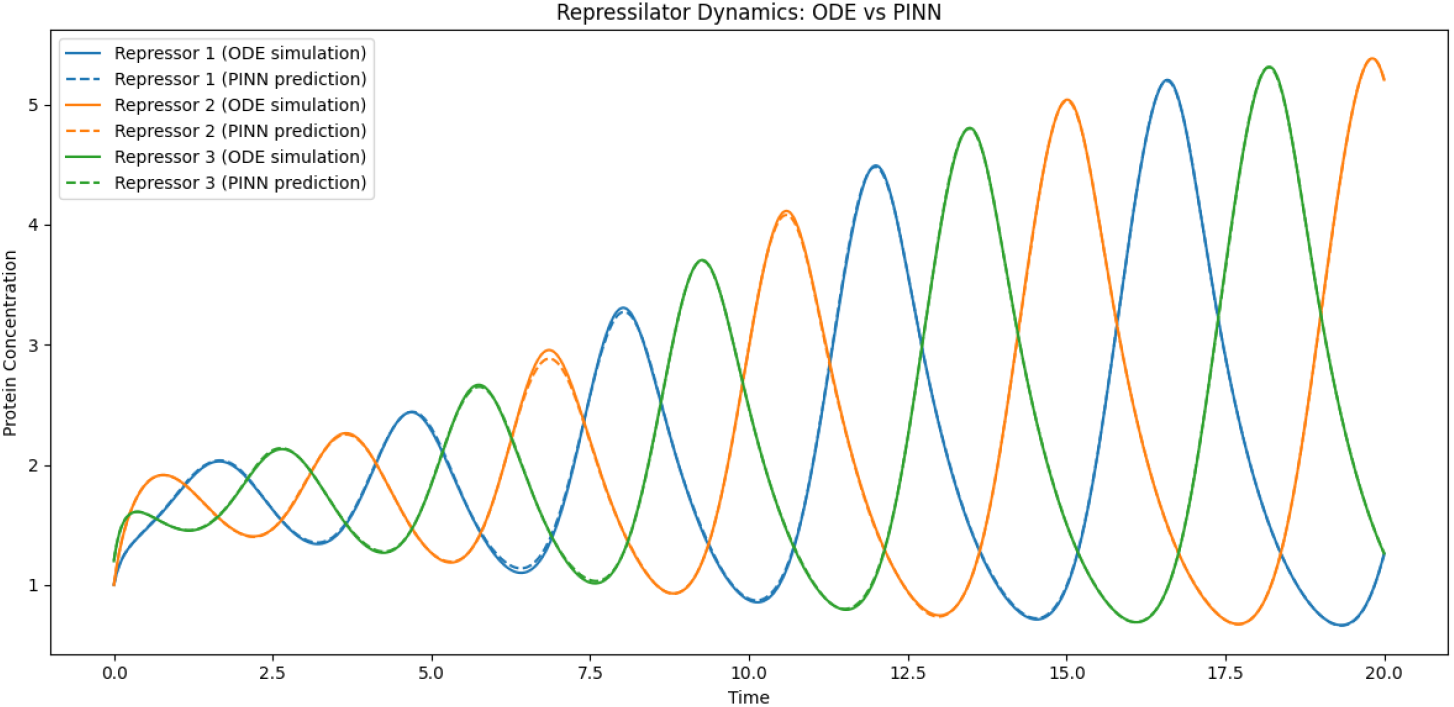
PINN reconstruction of an oscillatory repressilator trajectory. Solid lines are ODE simulations and dashed lines are PINN predictions.

### 3.2 Effect of Observation Noise

The noise-sweep experiment tests whether the physics residual compensates for corrupted measurements (Fig. 3). Performance degrades as the relative noise level increases from 1% to 10%. Parameter recovery remains comparatively stable over this range. The mean parameter error rises from 8.3 *×* 10^−3^ at 1% noise to 9.2 *×* 10^−3^ at 10% noise, with larger uncertainty at intermediate and high noise. State reconstruction responds more visibly to measurement corruption. The RMSE approximately doubles, increasing from 5.6 *×* 10^−3^ to 1.19 *×* 10^−2^ across the same sweep. The upward trend in both panels confirms that the inverse PINN benefits from clean observations even when the ODE residual regularizes the learned trajectory. At the same time, the small parameter errors in this controlled setting should not be read as general identifiability guarantees, because sparse sampling, hidden repressors, and unfavorable initial guesses can still produce biased estimates.

**Fig. 3.**
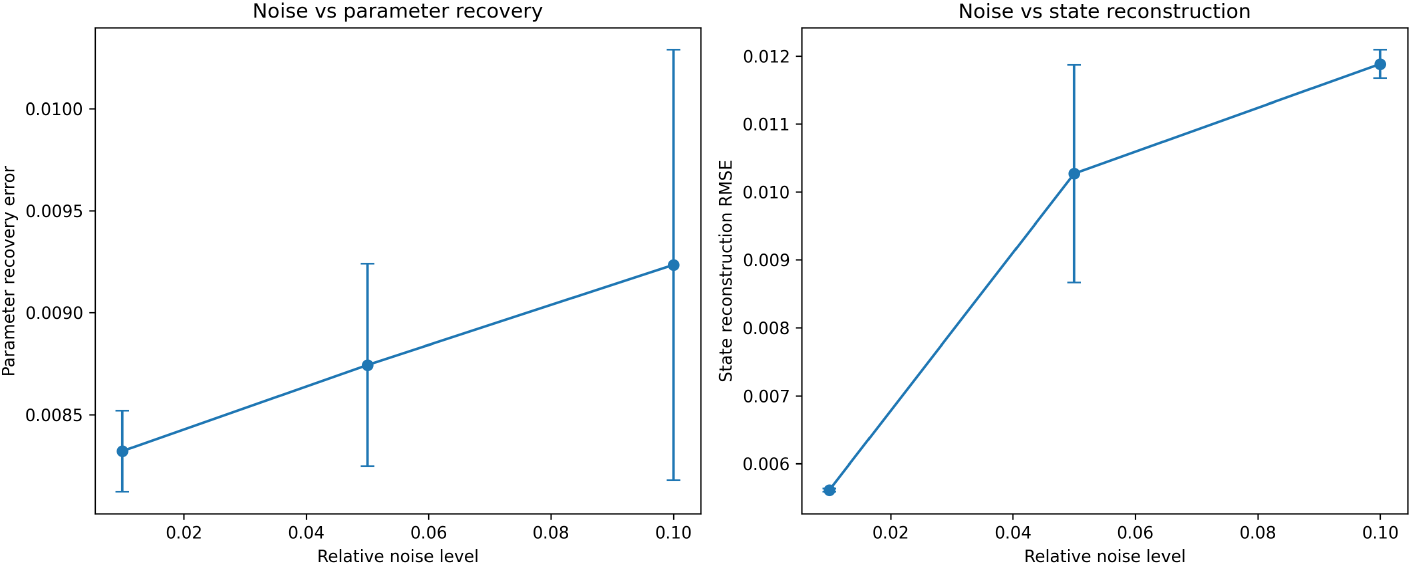
Effect of observation noise on inverse-PINN performance for the completed noise-sweep runs. The left panel reports the mean parameter recovery error, computed as the average relative error in *β* and *n*. The right panel reports the state-reconstruction RMSE against the clean simulated trajectory. Error bars show variability across the two repeated seeds.

### 3.3 Effect of Partial Observation

When all three repressors are observed, the inverse problem is best constrained because each equation contributes both a measured state and a measured regulator. Removing measurements makes the unobserved components latent variables whose values are inferred only through the ODE residual and the observed components. The two-repressor designs retain more information than the one-repressor design, but both increase variability across seeds. The one-repressor case is the most fragile because phase relationships among the repressors are only indirectly constrained.

### 3.4 Effect of Sampling Density

The sampling-density experiment shows the role of temporal coverage. Dense observations provide direct constraints on peaks, troughs, and phase lags. As the number of observation points decreases, the PINN can still interpolate smoothly because the dynamics are encoded in the residual, but parameter recovery becomes sensitive to whether the selected points cover informative regions of the oscillation. Very sparse sampling is therefore not merely a data-volume problem; it can remove the parts of the trajectory that distinguish different Hill coefficients.

### 3.5 Sensitivity to Initial Guesses

The initial-guess grid reveals optimization sensitivity. Starting close to the true values generally produces more reliable recovery, whereas distant initializations can converge to parameter values that reconstruct the observed trajectory acceptably but do not match the true parameters. This behavior is consistent with the non-convexity of neural-network training and with the practical nonidentifiability known in nonlinear biological calibration. Repeated seeds and multi-start strategies are therefore necessary when using inverse PINNs for parameter estimation.

### 3.6 STABLE Versus Oscillatory Regimes

The stable regime is easier for state reconstruction because trajectories converge to a fixed point. This same convergence can reduce parameter informativeness once the transient has decayed and many observations are concentrated near equilibrium. The oscillatory regime contains richer dynamical information, including amplitude and phase relations, but optimization is harder because errors in timing accumulate. In practice, the oscillatory regime is more useful for identifying parameters when sampling is adequate, while the stable regime is more forgiving for trajectory prediction.

## 4 Discussion

The experiments support the central expectation of the study. PINNs are effective at combining the repressilator equations with data and can reconstruct trajectories in both stable and oscillatory regimes. Their main value is not simply curve fitting; the ODE residual acts as a mechanistic regularizer that constrains latent states and discourages solutions that violate the known model.

At the same time, the study highlights a distinction between reconstruction and identification. A low trajectory RMSE does not guarantee accurate recovery of *β* and *n*. The Hill coefficient affects the nonlinearity of repression and can be weakly constrained when data are noisy, sparse, or missing some repressors. The behavior reflects practical identifiability limits rather than a PINN-specific failure, and it agrees with previous observations in biological parameter estimation [7, 9].

The partial-observation results are particularly relevant for biological experiments, where not every molecular species can be measured at every time point. A PINN can use the model to infer hidden components, but the inferred latent variables inherit the assumptions of the model. If the model structure is wrong, hidden-state reconstruction may look plausible while producing biased parameters. Methods designed to detect structural model deficiencies, such as topological augmentation [15], could complement inverse PINNs in this failure mode. A large physics residual that cannot be reduced by parameter adjustment may indicate that the assumed topology is itself the problem. Future work should therefore combine inverse PINNs with model-checking diagnostics of this kind, posterior or ensemble uncertainty, and experimental design criteria.

The repressilator is one instance of a finite catalogue of recurring regulatory topologies whose dynamical repertoire spans adaptation, pulse generation, bistability, and oscillation [13, 1]. Because the inverse PINN framework encodes model structure through the ODE residual, substituting a different motif topology mainly requires rewriting that residual while leaving the inference machinery unchanged. Each motif nevertheless imposes a distinct identifiability landscape. Negative autoregulation may be strongly constrained even by single-variable observations, whereas incoherent feed-forward loops can encode amplitude and duration in more separable parameters than the repressilator’s phase-coupled Hill coefficient. The genetic toggle switch illustrates a related challenge for bistable motifs, where deterministic structure and stochastic switching behavior may both matter for inference [16]. Systematic benchmarking of inverse PINNs across motif classes would therefore be more informative than single-system validation, providing a topology-indexed map of where this class of methods is reliable for reverse-engineering biological networks that can be decomposed into known motifs.

The initial-guess experiment also suggests that single-run PINN estimates should be treated cautiously. The optimization problem includes both neural-network weights and physical parameters, and the training loss can contain multiple basins. Multi-start training, parameter constraints, adaptive loss weighting, and reporting the distribution of estimates across seeds would make inverse PINN estimates more reliable for scientific use.

## 5 Conclusion

This paper presented an empirical characterization of inverse PINNs for parameter estimation in the repressilator model. The method can accurately reconstruct stable and oscillatory trajectories when the governing equations are correct, but parameter recovery is sensitive to data quality, observability, sampling, and initialization. The repressilator is a useful benchmark because it is small, interpretable, and dynamically nontrivial. The main conclusion is that PINNs are promising for reverse engineering ODE-based gene regulatory systems, provided that they are used with repeated runs, explicit identifiability checks, and observation designs that preserve phase and amplitude information.

